# A Hilbert-based method for processing respiratory timeseries

**DOI:** 10.1101/2020.09.30.321562

**Authors:** Samuel J. Harrison, Samuel Bianchi, Jakob Heinzle, Klaas Enno Stephan, Sandra Iglesias, Lars Kasper

## Abstract

In this technical note, we introduce a new method for estimating changes in respiratory volume per unit time (RVT) from respiratory bellows recordings. By using techniques from the electrophysiological literature, in particular the Hilbert transform, we show how we can better characterise breathing rhythms, with the goal of improving physiological noise correction in functional magnetic resonance imaging (fMRI). Specifically, our approach leads to a representation with higher time resolution and better captures atypical breathing events than current peak-based RVT estimators. Finally, we demonstrate that this leads to an increase in the amount of respiration-related variance removed from fMRI data when used as part of a typical preprocessing pipeline.

Our implementation will be publicly available as part of the PhysIO package, which is distributed as part of the open-source TAPAS toolbox (translationalneuromodeling.org/tapas).

**Highlights:**

- We introduce a new estimator for respiratory volume per unit time from respiratory recordings.
- We demonstrate how this is able to accurately characterise atypical breathing events.
- This removes significantly more variance when used as a confound regressor for fMRI data.
- Our implementation will be included in PhysIO, released as part of TAPAS: translationalneuromodeling.org/tapas.

## 1 Introduction

There has been much recent interest in the impact of global artefacts on fMRI data [Schölvinck et al. 2010; Satterthwaite et al. 2013; Burgess et al. 2016; Ciric et al. 2017; Murphy and Fox 2017; Liu et al. 2018], with many studies finding a link between physiological processes—particularly breathing—and these global signals [Power et al. 2017a; b; Byrge and Kennedy 2018; Power et al. 2019; 2020]. However, despite the fact that models for these physiological processes and their impact on fMRI are well established [Glover et al. 2000; Birn et al. 2006; 2008; Chang et al. 2009; Chang and Glover 2009; Murphy et al. 2013], much recent work has focused on improved data-driven methods for artefact removal [Glasser et al. 2018; Power et al. 2018; Aquino et al. 2020]. This is likely for two reasons. Firstly, even though physiologically-derived confounds have the advantage that they have a much greater a priori validity if one is concerned about mistakenly removing neural signal, there is a cost: they require extra data to be collected, inspected, and analysed [Glasser et al. 2018; 2019; Power 2019; Power et al. 2020]. Secondly, as Power et al. [2020] recently demonstrated, these fMRI artefacts typically arise in the context of unusual breathing events—very deep breaths, apnoeas, etc.—that are by definition hard for algorithms designed with ‘normal’ tidal breathing in mind to properly detect and characterise.

What we introduce here is a method, inspired by work from the electrophysiological literature, that seeks to address both of these concerns in the context of preprocessing recordings from respiratory bellows. We introduce a new estimator for respiratory volume per unit time (RVT) that does not require peak detection—a process that often requires post hoc manual intervention—and demonstrate that this accurately captures atypical breathing events. Empirically, when used as a confound regressor, this also removes more variance from fMRI data compared to a peak-based RVT estimator. Finally, for a much more detailed discussion of both breathing and breathing-related issues pertaining to fMRI than space affords in this technical note, we refer the reader to the excellent overview in Power et al. [ibid.].

### 1.1 Respiratory models and fMRI denoising

In the context of removing global fMRI confounds, the aim of the most commonly utilised breathing-related metric, RVT [Birn et al. 2006], is to collapse simple measurements of breathing—typically from peripheral physiological recordings, such as respiratory bellows—down to a metric that will correlate over time with the pressures of blood gases (e.g. pCO_2_ and pO_2_)^1^. There are two components to this: an increase in breathing rate *or* an increase in breathing depth will tend to, for example, decrease pCO_2_. As such, RVT is simply the product of respiratory volume (RV) and breathing rate.

The other component to these fMRI-based models is the mapping from RVT to the blood-oxygen-level-dependent (BOLD) contrast. Any changes in the pressures of blood gases impact on key constituents of the BOLD contrast, such as blood volume (via vasodilation) [Gauthier and Fan 2019]. To account for these physiological processes and the inherent delays in the circulatory system, a respiratory response function (RRF) [Birn et al. 2008] is used to map from RVT to the BOLD signal. This is directly analogous to the hæmodynamic response function which describes the mapping from neural activity to BOLD, and similar physiological models exist for changes in heart rate [Chang et al. 2009].

### 1.2 RVT estimators

As we describe in more detail later, current methods typically characterise the recordings from respiratory bellows via peak detection, but this has issues in terms of both robustness and temporal resolution. Here, we use a decomposition based on the Hilbert-transform to characterise the respiratory recordings; this is a technique that has been more widely used in the context of characterising complex oscillatory waveforms from electrophysiological data [Brookes et al. 2011; Hipp et al. 2012; Luckhoo et al. 2012; Engel et al. 2013; Voytek et al. 2013; Cole and Voytek 2017; Nguyen et al. 2019]. The crucial link is that the quantities that we need to define RVT, breathing depth and rate, are simply the amplitude and (instantaneous) frequency of the breathing-related oscillation in the bellows recordings.

By way of contrast, RVT is typically estimated via peak detection, as described by Birn et al. [2006]. Firstly, one detects points of maximum inhalation and exhalation (i.e. peaks and troughs in the respiratory waveform). Given these, respiratory volume is defined as the difference in belt positions between peaks and troughs, and breathing rate is the reciprocal of the time between successive peaks (with the relevant quantities linearly interpolated as necessary). However, peak detection is not robust to noisy recordings and fundamentally constrains the temporal resolution of the model to the observed breath durations. Taken together, these two issues are why RVT-based measures can often ‘miss’ deep breaths [Power et al. 2020].

As such, alternatives to RVT with better estimability have been proposed. For example, Chang et al. [2009] define a simple alternative to peak-based methods by simply taking the standard deviation of the respiratory signal in a small window around the timepoint of interest. While this has some of the purported benefits of our method, namely that it ‘does not rely on the accuracy of peak detection required for breath-to-breath computations’, it does not directly capture the effects of changes in breathing rate. Similarly, Power et al. [2018] propose what is essentially a hybrid of our approach and the above, by calculating the standard deviation of the respiratory envelope in a small window. Again, this is a peak-free method, but does not permit a clean dissociation of breathing depth and rate.

While the above two measures are more robust, they lack the direct physiological interpretability with regard to blood gas pressures that RVT offers. Therefore, our aim is to estimate RVT itself, and all comparisons in this technical note are with a peak-based RVT estimator. We use the implementation in the PhysIO package [Kasper et al. 2017], and this is publicly available as part of the TAPAS toolbox (translationalneuromodeling.org/tapas).

## 2 Methodology

### 2.1 Properties of the Hilbert transform and analytic signal

The Hilbert transform is central to our method as it allows us to derive an alternative representation of the respiratory recordings. For a given real-valued signal, *s*(*t*), we can define a unique analytic signal^2^, *s*_*a*_(*t*), in terms of the Hilbert transform ℋ [Gabor 1946]:

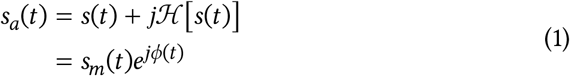

Where the second line is simply a rearrangement of the first into polar coordinates. Much more detail on the properties of this approach can be found in, for example, Boashash [1992a] and Huang et al. [2009].

The important aspect of this representation for the current work is the way we have split our signal into two components: an instantaneous magnitude, *s*_*m*_(*t*), and an instantaneous phase, *ϕ*(*t*). We can do one final rearrangement to make this more explicit:

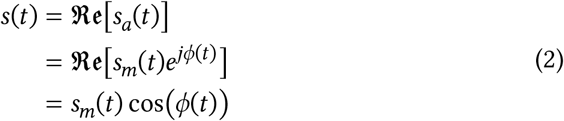

We plot this decomposition of a signal recorded from respiratory bellows in Figure 1. Note that we use the term amplitude envelope interchangeably with magnitude, as this is the common description in the electrophysiological literature.

**Figure 1:**
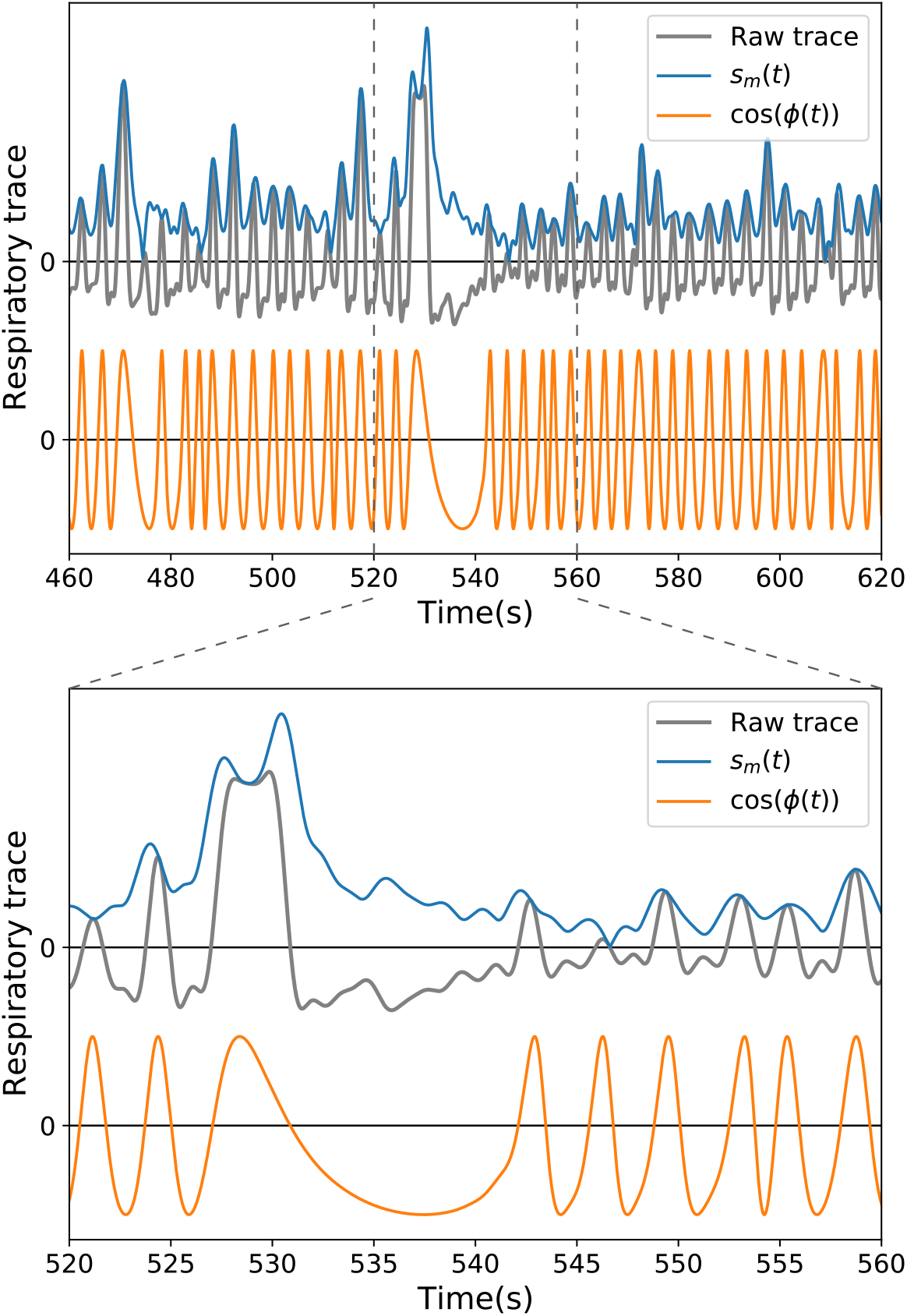
Example of the properties of the respiratory signal we extract via the Hilbert transform. The lower panel zooms in on a portion of the trace shown in the upper panel. Within each panel, in the upper half we overlay the amplitude envelope, *sm*(*t*), on the respiratory bellows trace. In the lower half we show the information carried by the instantaneous phase, which we illustrate as cos(*ϕ*(*t*)) as per Equation 2. The product of this oscillatory component and the amplitude recovers the original signal. A version of this plot showing a larger portion of the recorded signal is shown in Supplementary Figure S1.

Finally, these two terms are exactly what we need to calculate RVT. The respiratory volume (i.e. the difference between successive peaks and troughs) is simply twice the signal amplitude, *s*_*m*_(*t*). Similarly, the breathing rate is the instantaneous frequency, *s*_*f*_(*t*), which is the temporal derivative of the instantaneous phase:

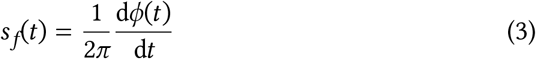

As a last step, we apply a low pass filter to these quantities to remove within-cycle fluctuations. The breathing rhythm is not sinusoidal, so the above terms will contain some high-frequency content that represents within-breath modulations of the waveform shape. Here, as in a typical preprocessing pipeline, RVT is convolved with the RRF—which is itself an aggressive low-pass filter—so this is not strictly necessary for fMRI denoising, but we do so because it affords more intuitive visualisations of the respiratory decomposition^3^.

### 2.2 Practical considerations and algorithmic approach

While the Hilbert transform is theoretically well suited to the problem at hand, deriving the necessary quantities can present a formidable practical challenge [Boashash 1992b; Huang et al. 1998]. Noise—or, more specifically, multiple local extrema that are not part of zero-centred oscillations—can cause the instantaneous frequency to become negative [Huang et al. 2009]. Correcting for these effects is a core part of the empirical mode decomposition (EMD) [Huang et al. 1998], whereby a single signal is decomposed into multiple modes, each of which is a monocomponent signal that admits a ‘well behaved’ decomposition into amplitude and phase terms. However, this is a data-driven approach that does not guarantee that the main breathing rhythm will be represented by a single mode.

As such, while we could take an EMD-based approach here, we do something simpler. Rather than decomposing the signal and then selecting the breathing-related modes post hoc, we assume that we can make an approximately mono-component signal by appropriately filtering the breathing signal. In practice, this requires a few rounds of iterative refinement, and the overall algorithm is detailed below. To aid the interpretation of the filtering operations below, note that the ‘normal’ breathing rate in adults—from both subjects resting in a supine position and undergoing MRI scanning—is approximately 0.25 Hz [Tobin et al. 1983; Power et al. 2019].

Our proposed algorithm for RVT estimation based on the Hilbert-transform consists of the following general steps:

1 | PhysIO preprocessing: Remove low frequency drifts (less than 0.01 Hz) from the breathing signal, and remove high-frequency noise above 2.0 Hz. This version of the breathing signal is used for both RETRO-ICOR and the peak-based RVT estimate in the physiological pipelines used in the Results section. This is the raw trace shown in Panel (a) of Figure 3.

**Figure 2:**
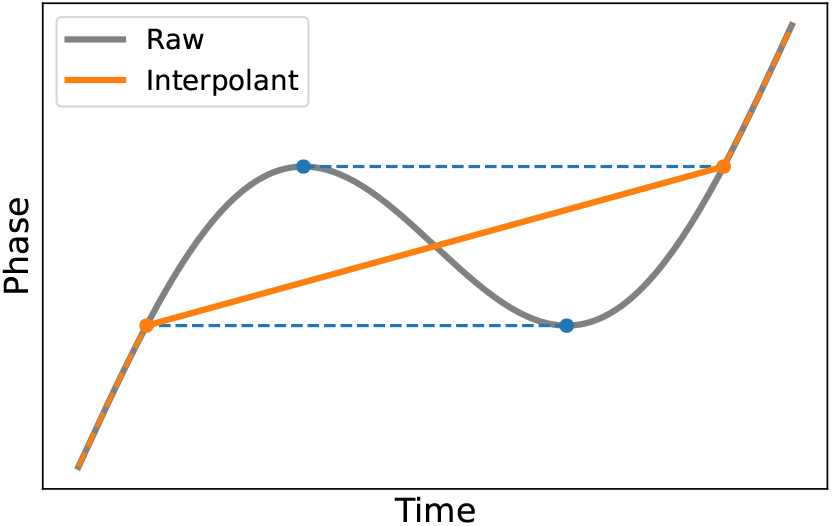
Graphical overview of the phase interpolation procedure used to ensure monotonicity. In brief, phase reversals are linearly interpolated from the time when the signal first crosses a threshold implied by the local minimum, to the equivalent time defined in terms of exceeding the local maximum.

**Figure 3:**
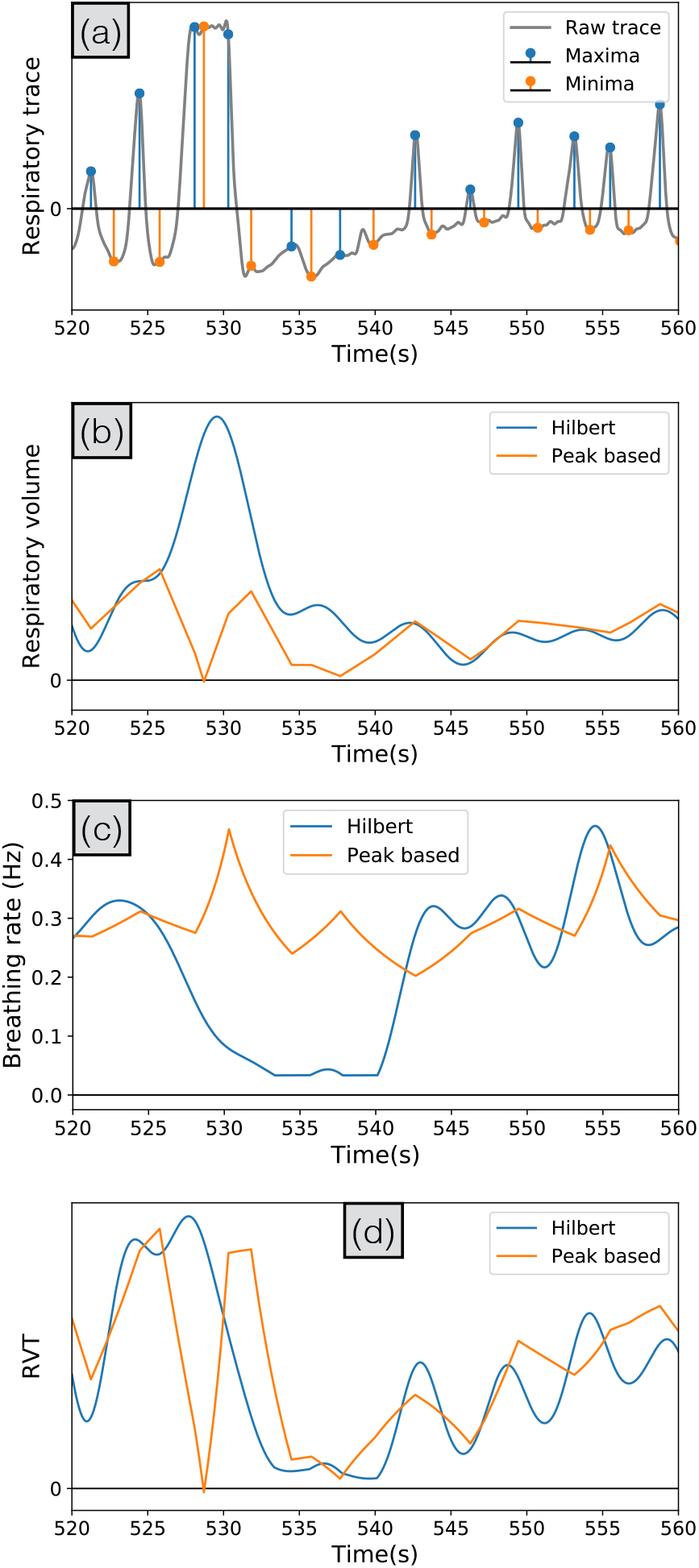
Comparison between the predictions from our Hilbert-based decomposition and a method based on peak detection. The portion of the respiratory trace is the same as in the lower panel of Figure 1, and includes a large inhalation at 530 s, followed by an apnoea lasting approximately 15 s. In **(a)** we show the detected peaks, which are the complete set of summary statistics used to define RVT as per Birn et al. [2006]. An extended version of this plot is shown in Supplementary Figure S3. In **(b), (c)**, and **(d)** we compare the inferred respiratory volume, breath durations, and RVT respectively.

2 | Lowpass filter the data again to more aggressively remove high-frequency noise above 0.75 Hz. This is the raw trace shown in Figure 1. The combined frequency response of these first two filtering steps is shown in Supplementary Figure S2.

3 | Decompose the signal into magnitude and phase components via the Hilbert transform.

4 | Linearly interpolate any periods where the phase timecourse decreases, using the procedure in Figure 2, to remove any artefactual negative frequencies. Reconstruct the oscillatory portion of the signal, cos(*ϕ*(*t*)), and lowpass filter at 0.75 Hz to remove any resulting artefacts. This procedure is repeated 10 times, with the new phase timecourse reestimated from the filtered oscillatory signal.

5 | Calculate RV from the signal magnitude and instantaneous breathing frequency as the numerical derivative of the phase timecourse. These estimated quantities are then filtered at 0.2 Hz to average the within-cycle fluctuations caused by the non-sinusoidal breathing waveform. Finally, estimates are thresholded to remove physiologically implausible values.

All filters, except for the common preprocessing, are 10^th^-order infinite impulse response, with the cut-offs stated in terms of the half-power frequency. Filters are run forwards and backwards to ensure zero-phase responses, and use 10 s of circular padding (i.e. `designfilt`, `padarray`, and `filtfilt` in MATLAB). The filters used during the preprocessing are 20^th^-order and use 100 s of padding due to the low cut-off frequency of the filter used to remove drifts in the signal.

## 3 Results

### 3.1 Data overview and preprocessing pipeline

Here, we demonstrate the performance of our method on data from a pharmacological fMRI study [Iglesias et al. 2021]. Briefly, we analyse data from 69 subjects who were scanned while performing an audio-visual associative learning task after receiving a minimal dose of either a dopaminergic antagonist (amisulpride), a cholinergic antagonist (biperiden), or placebo. Full details of the fMRI acquisition parameters, preprocessing pipeline, and task analyses can be found in the Supplementary Material.

We use this dataset to compare the performance of our RVT estimator to the peak-based approach implemented in PhysIO. As well as a set of qualitative comparisons, we run a group-level paired t-test to formally assess which method results in the stronger removal of respiratory-related variance.

### 3.2 Qualitative behaviour

In Figure 3 we compare the quantities inferred from the raw respiratory belt recordings. We focus on a single deep breath (i.e. a sigh [Li and Yackle 2017]), as this is the type of breath that Power et al. [2020] noted that RVT can miss.

In Panel (a) we show the detected maxima and minima, which amounts to the complete description of the data for peak-based methods (the equivalent Hilbert-based decomposition is shown in Figure 1). As detailed in Kasper et al. [2017], PhysIO uses a sophisticated peak detection algorithm that includes some assumptions about the regularity with which successive peaks appear in time. However, in the case of this particular complex waveform caused by the sigh, this results in local maxima and minima being incorrectly identified. This could be corrected post hoc—though this can be a time intensive procedure— or the assumption of regularity could be relaxed. However, as evidenced by other ambiguities in Supplementary Figure S3, and as discussed in Power et al. [2020], there is unlikely to be an optimal setting for peak detection algorithms. In this instance, one would manually mark a maximum at 528 s, a trough at 532 s, and the next maximum would be at 543 s; however, this is an incredibly sparse representation of 15 s of data and one which would conflate the increase in respiratory volume with the decrease in respiratory rate. Counter-intuitively therefore, the RVT method used here is more similar to an RV measure (i.e. the changes in breathing rate are small). Finally, note that while empirical results suggest that avoiding rate and depth changes cancelling in this manner may well improve denoising by reducing the number of ‘missed’ deep breaths, one would rather have an RVT estimator that was sensitive to these two effects separately [ibid.].

Finally, in Panels (b), (c), and (d) we compare the inferred respiratory volumes, breathing rates, and RVT itself. What is clear is that our method correctly identifies the changes in volume and rate, and furthermore, because we have a measure that is well defined for all points in time, these are not temporally overlapping. This leads to the expected changes in RVT, namely a rise driven by an increase in breathing volume, followed by a drop to well below baseline driven by the reduced breathing rate.

The Supplementary Material contains one extra visualisation of the data from this subject: in Supplementary Figure S4 we compare the inferred RVT regressors with the greyplot of the fMRI data itself.

### 3.3 Quantitative performance

In Figure 4 we compare the main effect of the RVT-based regressors at the group level. As detailed in the Supplementary Material, we run two separate group-level analyses to visualise the mean effect of RVT for the two different estimators, followed by a paired t-test analysing the differences between the methods. While both approaches show the expected pattern of strong positive effects distributed throughout the grey matter [Birn et al. 2006; 2008; Chang et al. 2009; Chang and Glover 2009; Power et al. 2017b], the bottom panel demonstrates how our method removes significantly more respiratory-related variance from the fMRI data.

**Figure 4:**
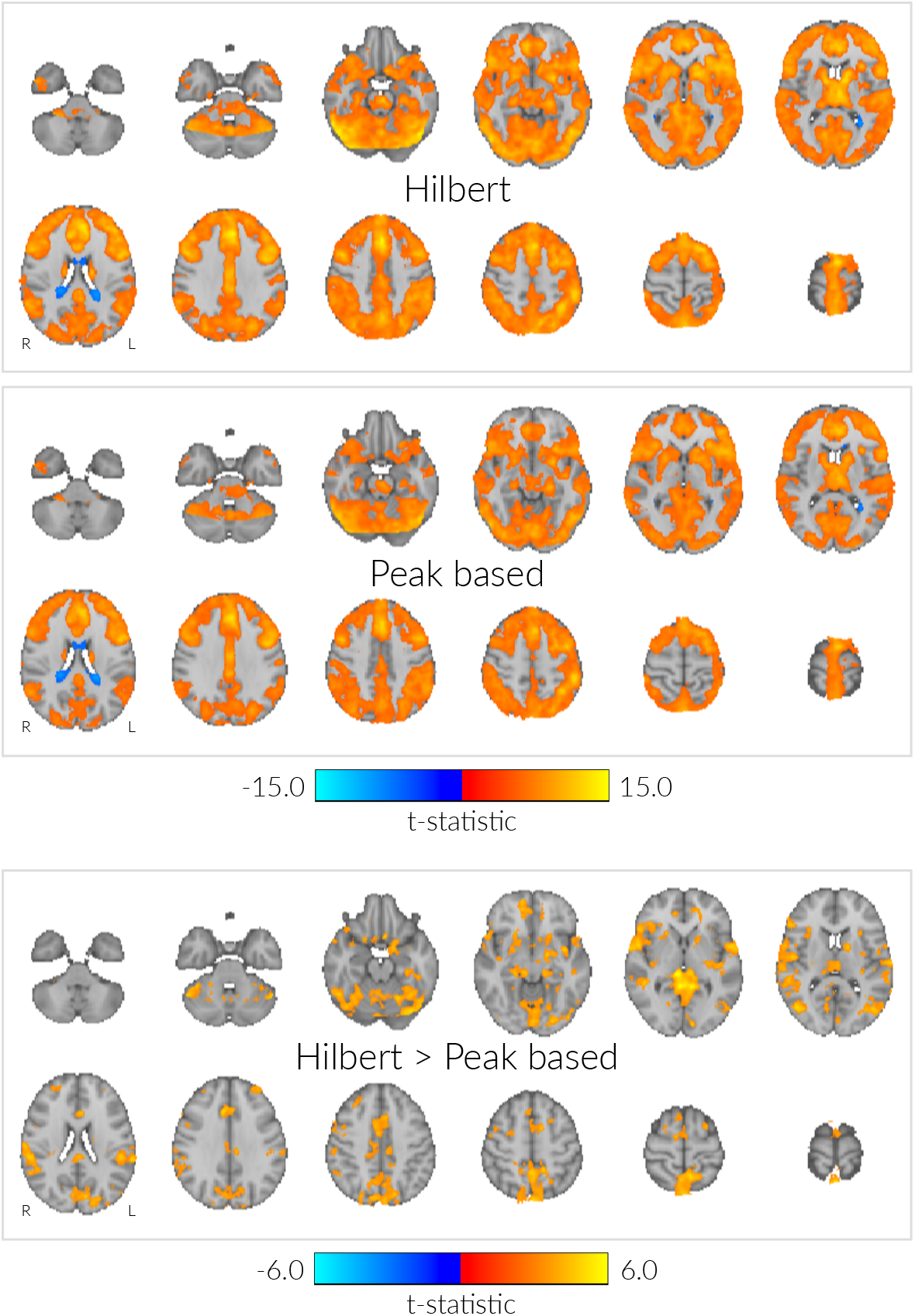
Comparison of the fMRI-RVT associations for our Hilbert-based RVT estimator and a method based on peak detection. In the upper two panels we show the t-stats for the main effect of the RVT regressors across the group (peak-level FWE-corrected at *p* = 0.05). In the lower panel we show the significant differences between the two as estimated via a group-level paired t-test (cluster-level FWE-corrected at *p* = 0.05 with cluster-forming threshold *p* = 0.001).

## 4 Discussion

In summary, we have demonstrated that the quantities we can derive via the Hilbert transform—instantaneous estimates of amplitude and phase—naturally map onto the quantities we need to estimate RVT. This furnishes us with a simple method that allows us to calculate a time-resolved version of RVT, rather than one defined in terms of a sparse set of peaks and troughs. We have demonstrated that this can better capture atypical breathing events, even in the context of non-sinusoidal oscillations with large and rapid changes in amplitude and phase. Finally, when convolved with the RRF as part of a typical fMRI preprocessing pipeline, our RVT estimates remove more respiratory-related variance from fMRI data than our baseline peak-detection-based method.

One of the aspects of our method we have emphasised throughout this technical note is the way it can dissociate changes in the depth and rate of breathing. However, the summary metric typically used for denoising, RVT, is the product of these two quantities. As Power et al. [2020] note, this introduces an ambiguity as ‘a larger-than-normal breath transpiring over a longer-than-normal time may appear quantitatively just like a typical breath occurring over a typical time’, or, in other words, RVT ‘is by definition relatively insensitive to outlier breath volumes so long as they [inversely] scale with breath times’. Our results speak to this in two ways. Firstly, what our results hint at is that this is primarily an artefact of the low time-resolution implied by reducing a signal down to its peaks alone, and not RVT itself. Figure 3 clearly shows how the changes in rate and depth we infer are not temporally coincident around a large and slow breath, and therefore we still see large changes in RVT in this instance. Secondly, however, if there are distinct differences in physiological responses to depth and rate changes [Birn et al. 2008; Lynch et al. 2020; Power et al. 2020] then our method provides both time series separately. This would allow one to derive fMRI regressors for both effects, which may improve the denoising performance for respiratory artefacts.

Next, in so far as our approach implies a model for the data, it is embodied by Equation 2: *s*(*t*) = *s*_*m*_(*t*) cos(*ϕ*(*t*)). There are two key assumptions here. Firstly, that the Hilbert transform provides a physically meaningful separation of fluctuations in magnitude and phase. The Hilbert transform is convenient, but splitting one time-varying signal into two time-varying quantities is an underdetermined problem that admits different solutions [Boashash 1992a]. Similarly, the interactions between the two quantities can themselves carry meaningful information [Huang et al. 2016; Nguyen et al. 2019]. Therefore, it may be that other decompositions or regularisation strategies improve performance. Secondly, the implicit assumption is that we can remove the vast majority of noise by appropriately filtering *s*(*t*). Again, this is an oversimplification in the presence of structured artefacts caused by, for example, contractions of the abdominal musculature unrelated to breathing or belt slippages [Power et al. 2020]. A more formal model might explain the data better here.

On the other hand, the simplicity of our model allows one to apply it robustly to a wide range of data, which may render it a useful quality control measure in and of itself. For example, given the magnitude and phase timecourses one could reconstruct the signal as modelled, and then compute a time-resolved measure of goodness-of-fit to the original recording. This may prove useful if it could, for example, be used to flag ‘bad’ timepoints that warrant manual inspection.

Finally, the other key aspect of the removal of these respiratory artefacts is the mapping from RVT to the BOLD signal via the RRF. It may be the case that our RVT estimator requires a subtly different RRF because it characterises large breaths in a slightly different manner. However, given the substantial variability in physiological responses over subjects and brain regions [Falahpour et al. 2013; Kassinopoulos and Mitsis 2019; Chen et al. 2020], we suspect that the use of flexibly parameterised, subject-specific response functions will dramatically improve the performance of denoising algorithms, and hope that the method proposed here is a useful addition to the overall physiological noise modelling pipeline.

## 5 Conclusion

In this technical note we have introduced a new method for estimating RVT from respiratory recordings for the purpose of removing physiological artefacts from fMRI data. We have demonstrated how this can both improve the detection of atypical breathing events and more strongly attenuate global fMRI artefacts as compared to current techniques.

## Acknowledgements

S.J.H. was supported by the grant #2017-403 of the Strategic Focal Area ‘Personalized Health and Related Technologies (PHRT)’ of the ETH Domain. K.E.S. was supported by the René and Susanne Braginsky Foundation and the University of Zurich. L.K. was supported by the NCCR ‘Neural Plasticity and Repair’ at ETH Zurich and the University of Zurich.

## Footnotes

1 This is distinct from and complementary to RETROICOR, which seeks to correct for instantaneous artefacts caused by the interaction between breathing-related motions and field changes, and cardiac pulsatility [Glover et al. 2000].

2 An analytic signal has no negative frequency components. While the Hilbert transform provides what is perhaps the simplest conceptual method for generating this (setting amplitudes of negative frequencies to zero, and doubling those for positive frequencies), there are infinite other ways of doing so.

3 For example, compare the double peak in the envelope at 530 s in Figure 1—caused by the square top to the waveform—with the filtered respiratory volume in Panel (b) of Figure 3.

## References

Aquino, K. M., Fulcher, B. D., Parkes, L., Sabaroedin, K. and Fornito, A. (2020). Identifying and removing widespread signal deflections from fMRI data: Rethinking the global signal regression problem. In: NeuroImage 212 (May 2020), p. 116614.

Birn, R. M., Diamond, J. B., Smith, M. A. and Bandettini, P. A. (2006). Separating respiratory-variation-related fluctuations from neuronal-activity-related fluctuations in fMRI. In: NeuroImage 31.4 (July 2006), pp. 1536–1548.

Birn, R. M., Smith, M. A., Jones, T. B. and Bandettini, P. A. (2008). The respiration response function: The temporal dynamics of fMRI signal fluctuations related to changes in respiration. In: NeuroImage 40.2 (Apr. 2008), pp. 644–654.

Boashash, B. (1992a). Estimating and interpreting the instantaneous frequency of a signal. I. Fundamentals. In: Proceedings of the IEEE 80.4 (Apr. 1992), pp. 520–538.

Boashash, B. (1992b). Estimating and interpreting the instantaneous frequency of a signal. II. Algorithms and applications. In: Proceedings of the IEEE 80.4 (Apr. 1992), pp. 540–568.

Brookes, M. J., Woolrich, M. W., Luckhoo, H., Price, D., Hale, J. R. et al.. (2011). Investigating the electrophysiological basis of resting state networks using magnetoencephalography. In: Proceedings of the National Academy of Sciences 108.40, pp. 16783–16788.

Burgess, G. C., Kandala, S., Nolan, D., Laumann, T. O., Power, J. D. et al.. (2016). Evaluation of Denoising Strategies to Address Motion-Correlated Artifacts in Resting-State Functional Magnetic Resonance Imaging Data from the Human Connectome Project. In: Brain Connectivity 6.9 (Nov. 2016), pp. 669–680.

Byrge, L. and Kennedy, D. P. (2018). Identifying and characterizing systematic temporally-lagged BOLD artifacts. In: NeuroImage 171 (May 2018), pp. 376–392.

Chang, C., Cunningham, J. P. and Glover, G. H. (2009). Influence of heart rate on the BOLD signal: The cardiac response function. In: NeuroImage 44.3 (Feb. 2009), pp. 857–869.

Chang, C. and Glover, G. H. (2009). Relationship between respiration, end-tidal CO2, and BOLD signals in resting-state fMRI. In: NeuroImage 47.4 (Oct. 2009), pp. 1381–1393.

Chen, J. E., Lewis, L. D., Chang, C., Tian, Q., Fultz, N. E. et al.. (2020). Resting-state “physiological networks”. In: NeuroImage 213 (June 2020), p. 116707.

Ciric, R., Wolf, D. H., Power, J. D., Roalf, D. R., Baum, G. L. et al.. (2017). Benchmarking of participant-level confound regression strategies for the control of motion artifact in studies of functional connectivity. In: NeuroImage 154 (July 2017), pp. 174–187.

Cole, S. R. and Voytek, B. (2017). Brain Oscillations and the Importance of Waveform Shape. In: Trends in Cognitive Sciences 21.2 (Feb. 2017), pp. 137–149.

Engel, A. K., Gerloff, C., Hilgetag, C. C. and Nolte, G. (2013). Intrinsic Coupling Modes: Multiscale Interactions in Ongoing Brain Activity. In: Neuron 80.4, pp. 867–886.

Falahpour, M., Refai, H. and Bodurka, J. (2013). Subject specific BOLD fMRI respiratory and cardiac response functions obtained from global signal. In: NeuroImage 72 (May 2013), pp. 252–264.

Gabor, D. (1946). Theory of communication. Part 1: The analysis of information. English. In: Journal of the Institution of Electrical Engineers - Part III: Radio and Communication Engineering 93 (26 Nov. 1946), pp. 429–44112.

Gauthier, C. J. and Fan, A. P. (2019). BOLD signal physiology: Models and applications. In: NeuroImage 187 (Feb. 2019), pp. 116–127.

Glasser, M. F., Coalson, T. S., Bijsterbosch, J. D., Harrison, S. J., Harms, M. P. et al.. (2018). Using temporal ICA to selectively remove global noise while preserving global signal in functional MRI data. In: NeuroImage 181 (Nov. 2018), pp. 692–717.

Glasser, M. F., (2019). Classification of temporal ICA components for separating global noise from fMRI data: Reply to Power. In: NeuroImage 197, pp. 435–438.

Glover, G. H., Li, T.-Q. and Ress, D. (2000). Image-based method for retrospective correction of physiological motion effects in fMRI: RETROICOR. In: Magnetic Resonance in Medicine 44.1 (July 2000), pp. 162–167.

Hipp, J. F., Hawellek, D. J., Corbetta, M., Siegel, M. and Engel, A. K. (2012). Large-scale cortical correlation structure of spontaneous oscillatory activity. In: Nature Neuroscience 15.6 (June 2012), pp. 884–890.

Huang, N. E., Hu, K., Yang, A. C. C., Chang, H.-C., Jia, D. et al.. (2016). On Holo-Hilbert spectral analysis: a full informational spectral representation for nonlinear and non-stationary data. In: Philosophical Transactions of the Royal Society A: Mathematical, Physical and Engineering Sciences 374.2065 (Apr. 2016), p. 20150206.

Huang, N. E., Shen, Z., Long, S. R., Wu, M. C., Shih, H. H. et al.. (1998). The empirical mode decomposition and the Hilbert spectrum for nonlinear and non-stationary time series analysis. In: Proceedings of the Royal Society of London A: Mathematical, Physical and Engineering Sciences 454.1971 (Mar. 1998), pp. 903–995.

Huang, N. E., Wu, Z., Long, S. R., Arnold, K. C., Chen, X. et al.. (2009). On Instantaneous Frequency. In: Advances in Adaptive Data Analysis 01.02, pp. 177–229.

Iglesias, S., Kasper, L., Harrison, S. J., Manka, R., Mathys, C. et al.. (2021). Cholinergic and dopaminergic effects on prediction error and uncertainty responses during sensory associative learning. In: NeuroImage 226 (Feb. 2021), p. 117590.

Kasper, L., Bollmann, S., Diaconescu, A. O., Hutton, C., Heinzle, J. et al.. (2017). The PhysIO Toolbox for Modeling Physiological Noise in fMRI Data. In: Journal of Neuroscience Methods 276 (Jan. 2017), pp. 56–72.

Kassinopoulos, M. and Mitsis, G. D. (2019). Identification of physiological response functions to correct for fluctuations in resting-state fMRI related to heart rate and respiration. In: NeuroImage 202 (Nov. 2019), p. 116150.

Li, P. and Yackle, K. (2017). Sighing. In: Current Biology 27.3, R88–R89.

Liu, X., Zwart, J. A. de, Schölvinck, M. L., Chang, C., Ye, F. Q. et al.. (2018). Subcortical evidence for a contribution of arousal to fMRI studies of brain activity. In: Nature Communications 9 (Jan. 2018), p. 395.

Luckhoo, H., Hale, J., Stokes, M., Nobre, A., Morris, P. et al.. (2012). Inferring task-related networks using independent component analysis in magnetoencephalography. In: NeuroImage 62.1, pp. 530–541.

Lynch, C. J., Silver, B. M., Dubin, M. J., Martin, A., Voss, H. U. et al.. (2020). Prevalent and sex-biased breathing patterns modify functional connectivity MRI in young adults. In: Nature Communications 11.1 (Oct. 2020), p. 5290.

Murphy, K., Birn, R. M. and Bandettini, P. A. (2013). Resting-state fMRI confounds and cleanup. In: NeuroImage 80 (Oct. 2013), pp. 349–359.

Murphy, K. and Fox, M. D. (2017). Towards a consensus regarding global signal regression for resting state functional connectivity MRI. In: NeuroImage 154 (July 2017), pp. 169–173.

Nguyen, K. T., Liang, W.-K., Lee, V., Chang, W.-S., Muggleton, N. G. et al.. (2019). Unraveling nonlinear electrophysiologic processes in the human visual system with full dimension spectral analysis. In: Scientific Reports 9.1 (Nov. 2019), p. 16919.

Power, J. D. (2019). Temporal ICA has not properly separated global fMRI signals: A comment on Glasser et al. (2018). In: NeuroImage 197 (Aug. 2019), pp. 650–651.

Power, J. D., Laumann, T. O., Plitt, M., Martin, A. and Petersen, S. E. (2017a). On Global fMRI Signals and Simulations. In: Trends in Cognitive Sciences 21.12 (Dec. 2017), pp. 911–913.

Power, J. D., Lynch, C. J., Dubin, M. J., Silver, B. M., Martin, A. et al.. (2020). Characteristics of respiratory measures in young adults scanned at rest, including systematic changes and “missed” deep breaths. In: NeuroImage 204 (Jan. 2020), p. 116234.

Power, J. D., Lynch, C. J., Silver, B. M., Dubin, M. J., Martin, A. et al.. (2019). Distinctions among real and apparent respiratory motions in human fMRI data. In: NeuroImage 201 (Nov. 2019), p. 116041.

Power, J. D., Plitt, M., Gotts, S. J., Kundu, P., Voon, V. et al.. (2018). Ridding fMRI data of motion-related influences: Removal of signals with distinct spatial and physical bases in multiecho data. In: Proceedings of the National Academy of Sciences 115.9, E2105–E2114.

Power, J. D., Plitt, M., Laumann, T. O. and Martin, A. (2017b). Sources and implications of whole-brain fMRI signals in humans. In: NeuroImage 146 (Feb. 2017), pp. 609– 625.

Satterthwaite, T. D., Elliott, M. A., Gerraty, R. T., Ruparel, K., Loughead, J. et al.. (2013). An improved framework for confound regression and filtering for control of motion artifact in the preprocessing of resting-state functional connectivity data. In: NeuroImage 64 (Jan. 2013), pp. 240–256.

Schölvinck, M. L., Maier, A., Ye, F. Q., Duyn, J. H. and Leopold, D. A. (2010). Neural basis of global resting-state fMRI activity. In: Proceedings of the National Academy of Sciences 107.22 (June 2010), pp. 10238–10243.

Tobin, M. J., Chadha, T. S., Jenouri, G., Birch, S. J., Gazeroglu, H. B. et al.. (1983). Breathing Patterns: 1. Normal Subjects. In: Chest 84.2 (Aug. 1983), pp. 202–205.

Voytek, B., D’Esposito, M., Crone, N. and Knight, R. T. (2013). A method for eventrelated phase/amplitude coupling. In: NeuroImage 64 (Jan. 2013), pp. 416–424.

