## Supplementary Material for "A Hilbert-based method for processing respiratory timeseries"

### A Hilbert-based method for processing respiratory timeseries: Supplementary Material

9th January 2021

#### 1 Methodology

In Supplementary Figure S1 we show an extended version of the decomposition of a respiratory bellows trace via the Hilbert transform, as per Figure 1 in the main text.

In Supplementary Figure S2 we plot the empirical frequency response of the combined preprocessing steps that we use to generate our approximately monocomponent breathing signal. In sum, the filters give a flat response between ≈0.01 Hz and ≈0.7 Hz, with strong attenuation (i.e. greater than −40 dB) outside of this range.

#### 2 Results

##### 2.1 Data overview and preprocessing pipeline

The full pharmacological fMRI study design, data, and analyses are described in Iglesias et al. [2021]. Drug administration was randomised and double-blinded, and scanning happened 90 minutes after the participants had received single oral dose of either a dopaminergic antagonist (400 mg amisulpride), a cholinergic antagonist (4 mg biperiden), or placebo. In this case, we analysed data from the audio-visual associative learning task, which is also described in Iglesias et al. [2013]. In total, 80 subjects were scanned while performing the task, but 11 subjects were excluded from the analyses here: 1 subject for missing electrocardiogram recordings, 5 subjects for too many missed trials or poor task performance, and 5 subjects for excessive motion (> 40 frames censored).

Analyses were run using MATLAB R2019a, SPM12 (r7487) [Friston et al. 2007] and the development version of PhysIO (branched from v7.1.0) [Kasper et al. 2017].

[a]Translational Neuromodeling Unit, University of Zurich & ETH Zurich, Zurich, Switzerland
[b]FMRIB, Wellcome Centre for Integrative Neuroimaging, University of Oxford, Oxford, UK
[c]Institute for Biomedical Engineering, ETH Zurich & University of Zurich, Zurich, Switzerland
[d]Wellcome Centre for Human Neuroimaging, University College London, London, UK
[e]Max Planck Institute for Metabolism Research, Cologne, Germany
[f]Techna Institute, University Health Network, Toronto, Canada

fMRI data were acquired on a 3 T Philips Achieva at a resolution of $2 \times 2 \times 3$ mm, and included 550 volumes at a TR of 2.5 s. fMRI data were preprocessed using SPM, including rigid-body motion correction, affine co-registration to the structural images, nonlinear registration from structural to MNI space, and smoothing (6 mm FWHM Gaussian kernel). First-level task regressors modelled the cue, button presses, targets (split into four regressors based on two factors: face/house stimuli presented and correctly/incorrectly predicted), and missed/invalid trials.

Physiological measures were collected via a breathing belt and 4-electrode electrocardiogram. An extensive set of physiological and motion regressors were estimated via PhysIO, including RETROICOR (model orders: cardiac phase: 3, respiratory phase: 4, interaction: 1) [Glover et al. 2000; Harvey et al. 2008], six motion parameters and their temporal derivatives [Friston et al. 1996], and spike regressors for motion censoring (framewise displacement > 0.5 mm) [Power et al. 2012; Satterthwaite et al. 2013]. RVT and heart rate were convolved with the RRF and CRF respectively [Birn et al. 2008; Chang et al. 2009], and lagged versions of each regressor (−5 s, 0 s, 5 s, 10 s) were included to account for variable latencies [Birn et al. 2006; Chang and Glover 2009].

At the single-subject level, all task, motion, and physiological regressors were entered into a GLM analysis. This was repeated twice: separately for both the Hilbert- and peak-based RVT estimators. The main effect of interest was the t-statistic from the average contrast over all four lagged RVT-based regressors.

These results were passed into two different group-level analyses. Firstly, for visualisation purposes, a simple group-level mean was calculated separately for each RVT estimator. Secondly, all results were fed into a paired t-test design to assess the statistical significance of any differences between methods. For all group-level analyses, drug, age, weight and two questionnaire-based sleepiness scales were included as confound regressors.

Finally, note that we did not observe any significant differences between the different RVT regressors when looking at the simple effects of task (i.e. face v. house and correct v. incorrect). The more complex model-based analyses of this task are presented in Iglesias et al. [2021].

#### 2.2 Qualitative behaviour

In Supplementary Figure S3 we show an extended portion of the respiratory trace, overlaid with the detected peaks. As well as the issues around the central deep breath discussed in the manuscript, there are clearly other areas of ambiguity (e.g. around 570 s and 610 s).

In Supplementary Figure S4 we compare the inferred respiratory regressors to a greyplot of the fMRI data for the exemplar subject used throughout the manuscript. There are several instances, for example at around 500 s, where the global decreases in the fMRI signal are better captured by the Hilbert-based regressor.

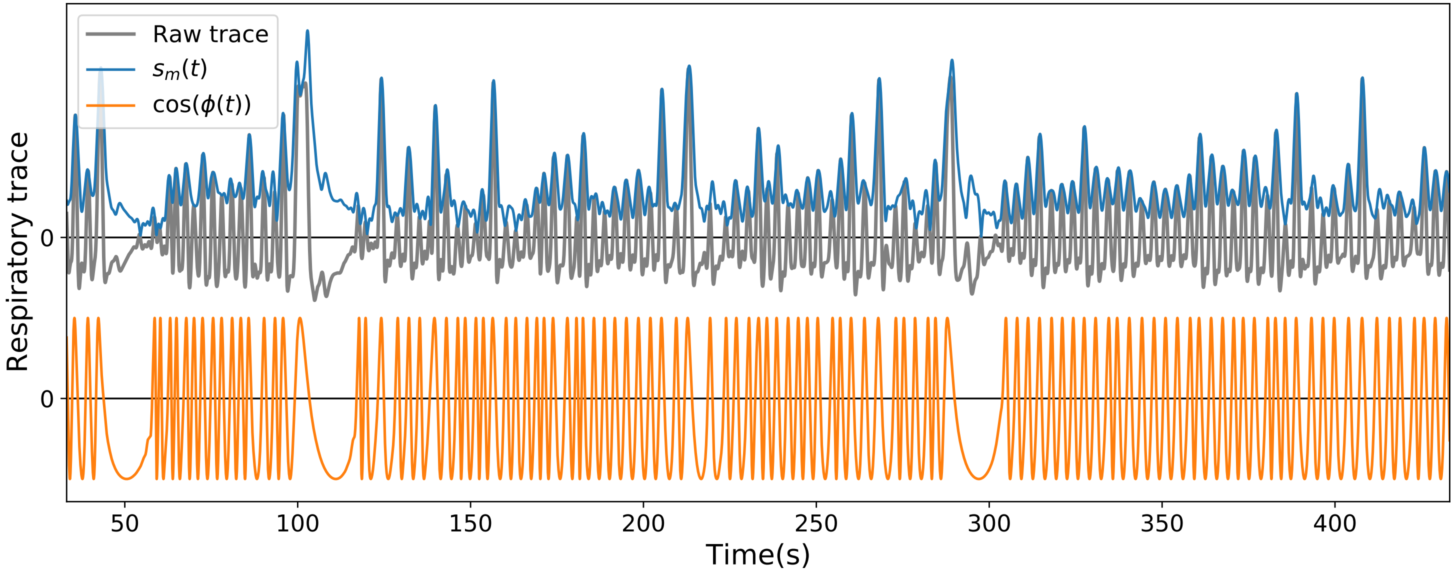


**Supplementary Figure S1:** Example of the properties of the respiratory signal we extract via the Hilbert transform. The plot is as per Figure 1 in the main text, but shows data from the same subject for a different and more extended period of time. In the upper half we overlay the amplitude envelope, $s_m(t)$, on the respiratory signal. In the lower half we show the information carried by the instantaneous phase, which we illustrate as $\cos(\phi(t))$. The product of this oscillatory component and the amplitude recovers the original signal.

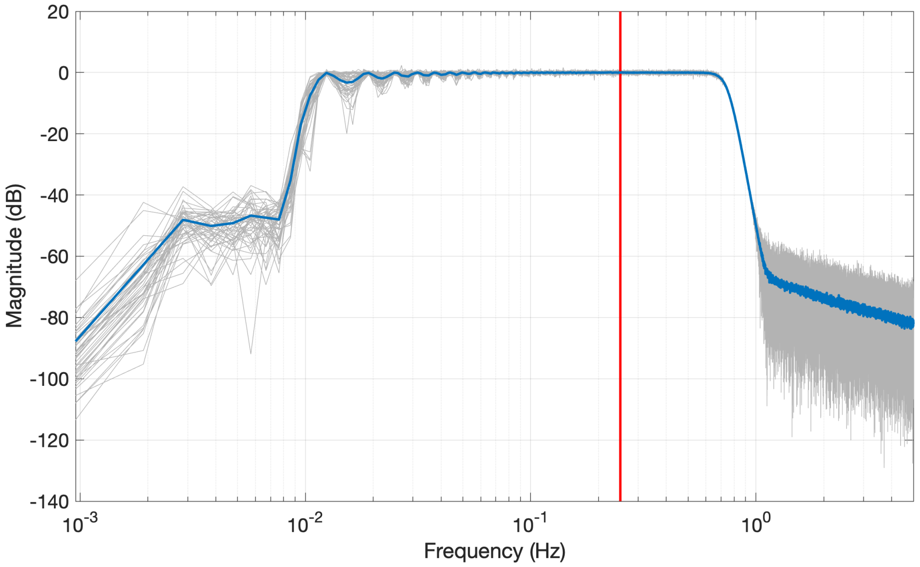


Supplementary Figure S2: Empirical frequency response of the preprocessing pipeline applied to the raw respiratory traces. As described in more detail in the main text, the pipeline detrends the raw data via a high-pass filter, and then removes noise via two low-pass filtering steps. The empirical transfer function is estimated via Welch's averaged periodogram method, as implemented in `tfestimate` in MATLAB. Each grey line represents the response to an input of white Gaussian noise at a sampling frequency of 250 Hz, with the blue line representing the mean over 50 realisations. The vertical red line indicates the typical breathing rate of 0.25 Hz mentioned in the main text.

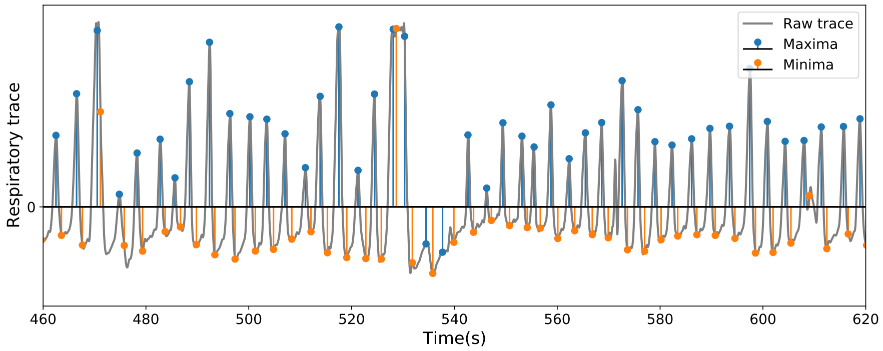


Supplementary Figure S3: Extended portion of the respiratory signal overlaid with detected peaks. The plot is as per Figure 3(a) in the main text, but shows data from the same subject for a more extended period of time. The data portion is the same as in the upper panel of Figure 1 in the manuscript.

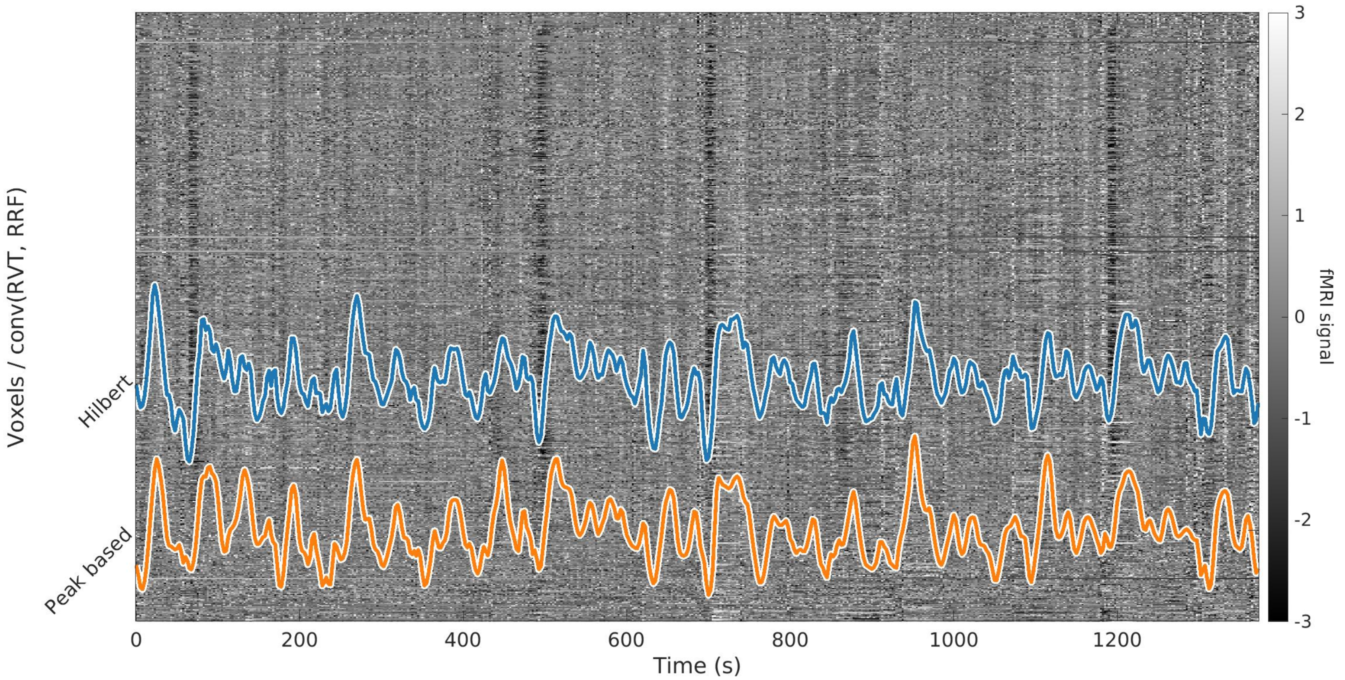


**Supplementary Figure S4:** Comparison of the RVT regressors (after convolution with the RRF) with the raw fMRI data. The data is from the same exemplar subject as the rest of the manuscript. For visualisation purposes, the RETROICOR and 12 motion regressors are removed from the detrended fMRI data, and the voxelwise time courses were z-scored.

#### References


Birn, R. M., Diamond, J. B., Smith, M. A. and Bandettini, P. A. (2006). *Separating respiratory-variation-related fluctuations from neuronal-activity-related fluctuations in fMRI*. In: *NeuroImage* 31.4 (July 2006), pp. 1536–1548.

Birn, R. M., Smith, M. A., Jones, T. B. and Bandettini, P. A. (2008). *The respiration response function: The temporal dynamics of fMRI signal fluctuations related to changes in respiration*. In: *NeuroImage* 40.2 (Apr. 2008), pp. 644–654.

Chang, C., Cunningham, J. P. and Glover, G. H. (2009). *Influence of heart rate on the BOLD signal: The cardiac response function*. In: *NeuroImage* 44.3 (Feb. 2009), pp. 857–869.

Chang, C. and Glover, G. H. (2009). *Relationship between respiration, end-tidal CO2, and BOLD signals in resting-state fMRI*. In: *NeuroImage* 47.4 (Oct. 2009), pp. 1381–1393.

Friston, K. J., Ashburner, J., Kiebel, S., Nichols, T. and Penny, W., eds. (2007). *Statistical Parametric Mapping: The Analysis of Functional Brain Images*. Academic Press.

Friston, K. J., Williams, S., Howard, R., Frackowiak, R. S. J. and Turner, R. (1996). *Movement-Related effects in fMRI time-series*. In: *Magnetic Resonance in Medicine* 35.3 (Mar. 1996), pp. 346–355.

Glover, G. H., Li, T.-Q. and Ress, D. (2000). *Image-based method for retrospective correction of physiological motion effects in fMRI: RETROICOR*. In: *Magnetic Resonance in Medicine* 44.1 (July 2000), pp. 162–167.

Harvey, A. K., Pattinson, K. T. S., Brooks, J. C. W., Mayhew, S. D., Jenkinson, M. et al. (2008). *Brainstem functional magnetic resonance imaging: Disentangling signal from physiological noise*. In: *Journal of Magnetic Resonance Imaging* 28.6, pp. 1337–1344.

Iglesias, S., Kasper, L., Harrison, S. J., Manka, R., Mathys, C. et al. (2021). *Cholinergic and dopaminergic effects on prediction error and uncertainty responses during sensory associative learning*. In: *NeuroImage* 226 (Feb. 2021), p. 117590.

Iglesias, S., Mathys, C., Brodersen, K. H., Kasper, L., Piccirelli, M. et al. (2013). *Hierarchical Prediction Errors in Midbrain and Basal Forebrain during Sensory Learning*. In: *Neuron* 80.2 (Oct. 2013), pp. 519–530.

Kasper, L., Bollmann, S., Diaconescu, A. O., Hutton, C., Heinzle, J. et al. (2017). *The PhysIO Toolbox for Modeling Physiological Noise in fMRI Data*. In: *Journal of Neuroscience Methods* 276 (Jan. 2017), pp. 56–72.

Power, J. D., Barnes, K. A., Snyder, A. Z., Schlaggar, B. L. and Petersen, S. E. (2012). *Spurious but systematic correlations in functional connectivity MRI networks arise from subject motion*. In: *NeuroImage* 59.3 (Feb. 2012), pp. 2142–2154.

Satterthwaite, T. D., Elliott, M. A., Gerraty, R. T., Ruparel, K., Loughead, J. et al. (2013). *An improved framework for confound regression and filtering for control of motion artifact in the preprocessing of resting-state functional connectivity data*. In: *NeuroImage* 64 (Jan. 2013), pp. 240–256.
